# Neural Circuit Mechanisms Underlying Dominance Traits and Social Competition

**DOI:** 10.1101/2024.12.04.626906

**Authors:** Han Yan, Jin Wang

## Abstract

The survival of animals often hinges on their dominance status, established through repeated social competitions. The dorsomedial prefrontal cortex (dmPFC) plays a pivotal role in regulating these competitions, yet the formation of intrinsic traits like grit and aggressiveness, crucial for competitive outcomes, remains poorly understood. In this study, we constructed a dmPFC circuit model based on experimental recordings to replicate the characteristic activities of dmPFC neurons during various behavioral patterns observed in the dominance tube test. Our findings reveal that the dmPFC circuit supports bistable behavior states—effortful and passive—depending on external conditions. This bistability is essential for understanding how animals adapt their behaviors in social competitions, thereby influencing the establishment of social hierarchies. Our results indicate that increased self-excitation in pyramidal neurons within the dmPFC enhances the robustness of effortful behaviors, akin to perseverance, but reduces flexibility in responding to rapid external changes. This suggests that dominance status benefits more from perseverance than from increased aggression. Additionally, our study shows that when rapid responses to external signals are necessary, the basal activity in dmPFC neurons can be reconfigured to enhance flexibility, albeit at higher energy costs. This research advances our understanding of the neural basis of social behavior and provides a framework for further exploration into how neural circuits contribute to complex behavioral traits, offering insights into the neural dynamics underlying social dominance. This research also opens avenues for investigating psychiatric and neurological disorders where these mechanisms may be disrupted.

## I. INSTRUCTION

In the animal kingdom, where the principle of “survival of the fittest” is paramount, social dominance is a critical factor^1–4^. Many species have evolved to live in structured groups where the establishment of a social hierarchy through repeated social competitions is essential for determining an individual’s access to crucial resources like food, mates, and shelter^3,5–8^. This access, in turn, significantly influences their physical and mental well-being. The formation of social hierarchies within group-living species is not solely a product of physical attributes like strength or size but also involves mental factors such as courage, persistence, and prior winning experiences. These factors, mediated by cortical brain regions that govern high cognitive functions, are crucial in dictating the outcomes of social competitions and the resultant social hierarchies^8–14^.

To explore the neural underpinnings of these processes, researchers have employed the dominance tube test, a method designed to assess social hierarchy by observing direct competitive interactions^15–17^. Recent research has highlighted the significant role of the dorsomedial prefrontal cortex (dmPFC) in influencing the outcomes of these competitions and thereby affecting the establishment of social hierarchies^16,18–20^. Additionally, studies have shown that synaptic plasticity within the mediodorsal thalamus-dmPFC pathway is crucial for encoding the “winner effect”, which increases the probability of future victories and reinforces the dominance hierarchy^21,22^. While the involvement of local microcircuits within the dmPFC in controlling social competition has been suggested^23^, the extent to which these circuits encode intrinsic traits or mental strengths remains to be fully elucidated.

Survival in animals demands the optimization of rewards while minimizing energy expenditure^24^. During social competitions, animals are required to select the most suitable behavioral response after assessing the cost-benefit ratio^25,26^. The neural circuitry involved operates in a challenging, noisy environment^27–30^, requiring the generation and maintenance of robust behavioral outcomes in response to specific sensory signals while accommodating input variations. If effortful behaviors, such as pushing and resistance in the tube test, fail to persist against these fluctuations and quickly transition to passive behaviors, such as stillness and retreat, the invested efforts yield no return^22,23^. Conversely, neuronal circuits must also display sufficient flexibility to adapt behaviorally to changing circumstances. These considerations give rise to questions regarding the ability of the circuit function in social competition to balance robustness and flexibility.

Our study aims to refine the understanding of how specific cognitive processes, often simplified as components of complex personality traits, are manifested in neural activity during social interactions. By doing so, we hope to contribute to the broader discourse on the neurobiological basis of social behavior and personality in animals. We constructed a dmPFC circuit model based on previous experimental recordings^22,23^. We developed a dmPFC circuit model grounded in prior experimental recordings. This model effectively captures the distinct activities of various dmPFC neuron types during different behavioral patterns observed in the tube test. Notably, our analysis indicates that these characteristic neural activities are only apparent under certain control parameter settings, leading to the formation of a bistable parameter region where both effortful and passive behaviors can be supported. Our findings suggest that the trait of perseverance can be enhanced within this circuit through increased self-excitation in pyramidal neurons, promoting robust effortful behaviors. However, this circuit configuration also impacts the system’s ability to swiftly adapt to external changes, a phenomenon we refer to as the robust-flexibility tradeoff. We propose that adjusting the system closer to the critical point, where the dynamical attractors associated with effortful and passive states are equally probable, could improve responsiveness to external stimuli, albeit at a higher energy cost. The specific architecture of this neural circuit, combined with dynamic adjustments based on environmental cues, plays a crucial role in determining competitive outcomes and establishing stable social hierarchies. In Section II.A, we describe the role of the dmPFC microcircuit in controlling social competition outcomes. Section II.B discusses the implications of our findings on how innate traits in social competition are shaped in the local dmPFC circuit, while Section II.C explores the thermodynamic costs associated with maintaining specific neural states and their impact on behavioral flexibility. Finally, Section III synthesizes our results and discusses their broader implications for understanding and manipulating social behavior in animals.

## II. RESULTS

### A. The role of the dmPFC microcircuit in controlling social competition outcomes

An animal’s social hierarchy is not solely determined by physical attributes such as body size or strength, but also influenced by intrinsic personality traits and past winning experiences, all regulated by higher cortical functions^8,10^. To unravel the neural mechanism underlying the interplay of intrinsic and extrinsic factors in determining social status, researchers have employed laboratory mice and devised methods to measure dominance hierarchy, such as the tube test. In this test, two mice compete for passage through a narrow tube, exhibiting effortful behaviors (push initiation, pushback, and resistance) or passive behaviors (stillness and retreat) when confronted by their opponents^22^. Winner mice demonstrate a higher frequency of pushes and resistance, and a lower frequency of retreats compared to loser mice^22,23^. Through the utilization of the dominance tube test, a specific neural population in the dorsomedial prefrontal cortex (dmPFC) has been identified as a key regulator in determining social dominance^16,18,19^. Notably, activation of dmPFC neurons, particularly the excitatory pyramidal (PYR) neurons, during the tube test elicits winning behaviors^16^.

Recent studies have shed light on the involvement of dmPFC interneurons in social behaviors^23,31–33^. Experimental recordings have demonstrated that inhibition of parvalbumin (PV) interneurons in the dmPFC promotes winning behaviors, while activation of PV interneurons induces losing behaviors^23^. Based on these findings, a microcircuit within the dmPFC has been proposed (Fig. 1(a)) to elucidate how local microcircuits in the medial prefrontal cortex (mPFC) control the outcomes observed in dominance tube test performance^23^. PV interneurons exhibit relatively high basal activity and exert mild tonic inhibition on PYR neurons. During retreats, PV neurons are activated, further suppressing PYR neurons. In push epochs, the experimental increase in PYR activity is hypothesized to arise from the disinhibitory effects resulting from the transient inhibition of PV neurons. However, intriguingly, PV neurons also show a significant increase in activity during push epochs, which contradicts the inhibitory circuit architecture. Additionally, behavioral patterns such as resistance without locomotion and stillness have been observed in the tube test^22,23^. The underlying mechanisms through which these behaviors emerge within the mPFC local circuitry and whether they can be attributed to specific circuit outputs involving distinct types of neurons remain unclear.

**FIG. 1.**
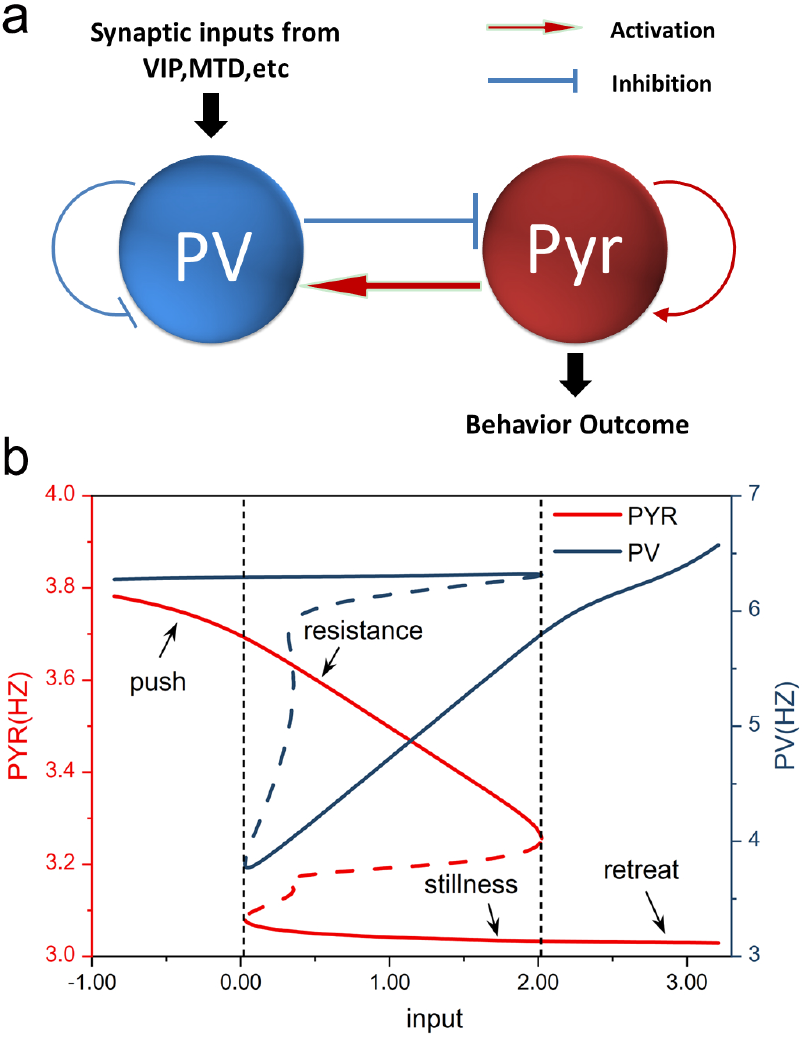
(a) The schematic diagram of the dmPFC circuit model. Excitatory pyramidal (PYR) neurons excite themselves and also parvalbumin (PV) interneurons. PV neurons, conversely, inhibit the activity of PYR neurons. (b) The phase diagram of the dmPFC circuit model with varied external inputs. The stable states on the lower branch of the solid curves correspond to the “passive” behaviors of mice in the tube tests. While, the stable states on the upper branch of the solid curves correspond to the “effortful” behaviors in the tube test. The dashed lines indicate bifurcation points that separate monostable and bi-stable regions. There is an increase/decrease in the activity of PYR neurons during the push/retreat epochs. PV neurons showed a significant increase in both push and retreat epochs. The effortful behaviors of the resistance can be identified through the characteristic activity in the bi-stable region.

To address these questions, we developed a local PYR-PV circuit model. Our circuit model is grounded in electrophysiological data from previous studies on dmPFC neuronal activity during social competition tasks^22,23^. We used approximate mean firing rates of PYR and PV neurons during different behavioral states to calibrate our model parameters. Detailed information on data sources and model construction is provided in the Appendix. Our model incorporates nonlinear terms that describe the interactions among PV and PYR neurons, including a self-excitation mechanism in the PYR population. This self-excitation is not to be interpreted as a literal self-stimulation of individual PYR neurons, but rather as a representation of the network dynamics within the dmPFC, where recurrent excitation among PYR neurons and inputs from other excitatory neurons can lead to sustained or enhanced activity levels. In the brain, such dynamics could arise from local circuits that include recurrent collaterals of PYR neurons or from long-range excitatory projections from other brain areas that synapse onto PYR neurons, contributing to their overall level of activity. Stimulus inputs carrying relevant sensory information from upstream brain regions, which play a role in determining the outcome of social competition, were modeled as inputs to PV neurons in our model. In the context of the tube test, the higher the rank of the opponent faced, the greater the input received by PV neurons. By exploring the parameter space of the model, including stimulus input strength and the connection strength of self-excitation in the PYR neural population, we discovered that distinct patterns of dmPFC neuronal activity observed in mice during the social dominance tube test could be replicated within bistable regions.

Fig. 1(b) illustrates the phase diagram, where the stimulus inputs to PV neurons were systematically varied while keeping all other parameters constant. Mathematically, the solid curves in the phase diagram represent the stable states of the dynamical system. The lower branch of the solid curves corresponds to stable states characterized by increased activity in PV neurons and inhibited PYR neurons, aligning with the “passive” behaviors displayed by mice in the tube test. In contrast, the upper branch of the solid curves represents stable states with heightened activity in both PV and PYR neurons, reflecting “effortful” behaviors in the tube test^23^. The push and retreat actions observed in the tube test can be identified as stable states located in the left and right regions of the phase diagram in Fig. 1(b), respectively. The dashed lines represent bifurcation points that separate mono-stable and bi-stable regions. We hypothesize that the effortful behavior of resistance occurs in the intermediate region of the phase diagram where bistability is present. This hypothesis finds support in electrophysiological recordings showing that the mean firing rate of PYR neurons during the resistance epoch is lower than during the push epoch^22^. Consistently, our model also reveals that the mean activity level of PYR neurons on the upper branch is relatively lower within the bistable region compared to the monostable region associated with push behaviors. In the bistable region, there exist two alternative stable states, effortful and passive responses, on the upper and lower branches, respectively. In realistic neural circuits with finite noise, initial states on the upper branch may occasionally transition to the lower branch, posing a challenge to maintaining robust effortful behavioral outputs within this bistable region compared to the monostable region. Consequently, the average activity of PYR neurons, which is read out by downstream brain regions responsible for executing dominance behavior, decreases. Furthermore, the behavioral pattern of “stillness” observed in the tube test is likely to occur around the bifurcation point that separates the right monostable region from the middle bistable region. States on the lower branch correspond to PYR activity states that are less inhibited compared to the retreat epoch, and the system around this region can robustly support passive behavioral outputs that are unlikely to transition to effortful behaviors on the upper branch.

### B. Innate traits in social competition are shaped in the local dmPFC circuit

In the tube test, a mouse transitions from a state of “stillness” to various behaviors as it encounters an opponent. The outcome of social competition is influenced by both external factors, such as the opponent’s state and prior history, and internal factors, including personality traits regulated by the local dmPFC circuit^8^. To understand how this local circuit influences dominance-related traits, we systematically manipulated key control parameters, such as the strength of self-excitation in PYR neurons, and analyzed the resulting circuit outputs based on its activity states. In our model, the strength of self-excitation in PYR neurons is quantified by a term comprising a constant part represented by the control parameter *a* and a normalized nonlinear function dependent on the mean activity of the PYR neural population. Figs. 2(a) and (b) illustrate phase diagrams with varying external inputs for the control parameter *a* = 26.4 and *a* = 28, respectively.

**FIG. 2.**
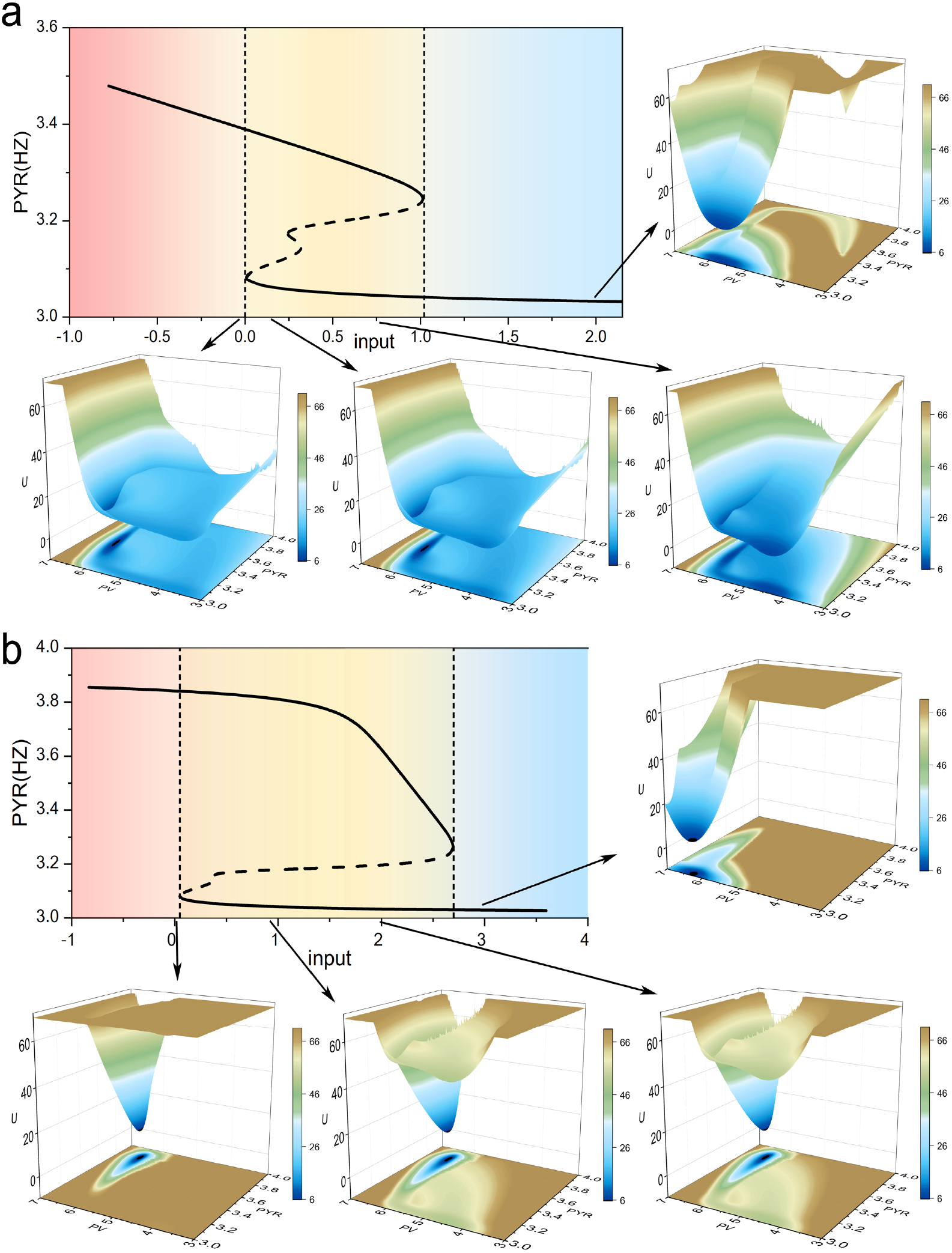
The phase diagram of the dmPFC circuit model and underlying attractor landscapes for varied external inputs with the strength of self-excitation in PYR neurons *a* = 26.8(subfigure (a)) and *a* = 28(subfigure (b)), respectively. The mono-stable regions supporting push and retreat behaviors are indicated by the red and blue colors, respectively. The bi-stable region supporting both effortful and passive behaviors is indicated by the yellow color. With strengthened self-excitation in PYR neurons(subfigure (b)), the stable states corresponding the effortful behaviors are much robust to changes in the external inputs.

As the self-excitation increases, the bi-stable phase expands into the original blue region, where retreats typically occur. This expansion suggests that the system becomes less likely to transition from the upper branch to the lower branch. Consequently, effortful behaviors become more resilient to increased external inputs, which represent opponents with higher social status, and transitions to passive retreats become less probable. Thus, the innate trait of perseverance in social competition, operationalized in our model as maintaining effortful behavior despite external pressures to retreat, can be fostered through heightened self-excitation in PYR neurons. This increased perseverance intuitively leads to more victories in social competition and, subsequently, a higher social status. Despite these changes, the red region corresponding to effortful pushes undergoes minimal change with increased self-excitation in PYR neurons, indicating that heightened self-excitation does not significantly enhance the probability of initiating assertive pushes in the local circuit, but rather increases the persistence of effortful behaviors like resistance. This theoretical result aligns with previous experimental findings suggesting that dominant animals do not necessarily exhibit more aggressive behaviors in broader contexts (e.g., attacks, chases)^22,34,35^, and our model suggests that increased dominance can be achieved through enhanced persistence in effortful behaviors rather than increased assertive pushes.

In the bi-stable region, both effortful and passive responses are supported. To gain a deeper understanding of how the local circuit influences behavioral outcomes in social competition, and particularly whether the expanded bi-stable region resulting from increased self-excitation in PYR also promotes more effortful behaviors in addition to reducing passive retreats, it is crucial to quantify the preference between the two alternatives within the bi-stable region. Quantification of the underlying dynamical attractor landscapes allows for a comprehensive analysis of noisy neural systems, encompassing the global stability of functional states and transitions between them (see the Appendix for detailed information)^36–39^. Fig. 2 illustrates the underlying attractor landscapes with varied external inputs. As the external inputs decrease, the dynamical attractors associated with passive behaviors (characterized by lower activity in PYR neurons) become destabilized, while the attractors associated with effortful behaviors (characterized by higher activity in PYR neurons) emerge and dominate. The local dmPFC circuit determines behavioral outcomes based on changes in the external environment, such as the state of the opponent. For a specific external input value (e.g., *input* = 1.8), the attractor for effortful behavior disappears in the circuit with lower self-excitation in PYR (*a* = 26.4), whereas the circuit with higher self-excitation in PYR (*a* = 28) can still sustain robust effortful behaviors. In other words, as the self-excitation in PYR increases, the attractors associated with effortful behaviors become more stable and less susceptible to shifts caused by varying external inputs. This stability is also quantitatively shown by the increased area under the curve in the phase diagrams, which indicates a larger parameter space where the effortful state is maintained despite fluctuations in external inputs. The distinct synaptic connections within the circuit give rise to different intrinsic traits in social competition.

In a specific range of external inputs, characterized by the coexistence of two alternative attractor states, the local circuit exhibits the ability to switch between effortful and passive behaviors under the influence of noise. This neurobiological phenomenon provides a potential foundation for characteristic performances observed in social competition, such as courage and fear. Importantly, the transition from one attractor state to the other requires the system to overcome a potential barrier within the bi-stable region. To assess the robustness of effortful and passive behaviors under different control parameters, we quantified the barrier heights between the corresponding attractors and the average transition time between them.

By examining three different values of self-excitation in PYR neurons, we discovered that increased external inputs to PV neurons decreased both the barrier height separating the effortful-behavior attractor from the passive-behavior attractor and the corresponding transition time (Fig. 3(a) and 3(c)). However, transitioning from passive behaviors to effortful behaviors took longer when crossing higher barriers with increased external inputs (Fig. 3(b) and 3(d)). Notably, both the average transition time from effortful to passive states and vice versa increased as self-excitation in PYR neurons was strengthened (indicated by the red dots compared to the blue ones). These results indicate that effortful behaviors become more resilient not only against increased excitatory inputs to PV neurons but also against the intrinsic fluctuations inherent in neural circuits with higher self-excitation in PYR.

**FIG. 3.**
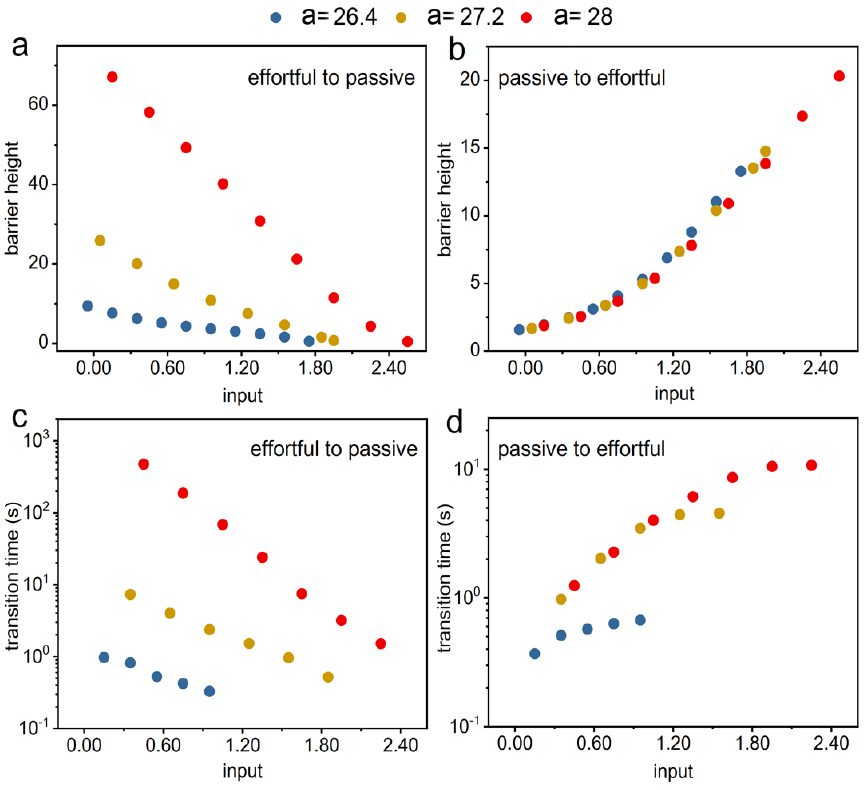
Robustness of the two alternative attractor states in the bistable region in terms of the transition time from one to the other and the barrier height in between inferred from the underlying landscape topography. For a specific value of the strength of the self-excitation in PYR, the decreased input to PV neurons induces increased transition time from the effort states to the passive states by crossing the higher barriers((a) and (c)). On the contrary, the decreased inputs to the PV neurons make it easier to shift to the effort states from the passive states((b) and (d)). For strengthened self-excitation in PYR indicated by different colors, the average transition time to passive behaviors are significantly increased with also slight increase in the transition time to the effortful states.

From an intuitive perspective, increased dominance in social competition appears to be facilitated by both the personality traits of perseverance and aggressiveness. However, our findings from Fig. 3c and 3d suggest that the perseverance resulting from increased self-excitation in PYR neurons primarily reduces the transitions from effortful behaviors to passive behaviors, without significantly increasing the transitions from passive behaviors to effortful behaviors within the bi-stable region. This implies that the level of basal aggression may not be heightened, aligning with experimental observations^22^. Therefore, our results suggest that animals displaying perseverance in competitions may also exhibit prudence, and an increase in social rank during competition may be attributed more to the innate trait of perseverance rather than heightened aggression.

Our model highlights the importance of perseverance, as indicated by the increased resilience of effortful behaviors within the bi-stable region of the dmPFC circuit. However, it is crucial to consider the context in which these behaviors occur. The tube test data, which underpins our model, captures a scenario where aggression is mild and typically involves familiar individuals. In more complex social structures or during the initial establishment of social hierarchies among unfamiliar animals, the dmPFC circuit might play a different role, potentially facilitating higher levels of aggression as a means to achieve dominance^40^. Therefore, while our findings underscore the significance of perseverance over aggression in the context of the tube test, they may not fully extrapolate to all forms of social competition. In light of these considerations, future work should aim to integrate data from a broader range of social interactions, including those involving unfamiliar individuals and higher intensity aggression, to develop a more comprehensive understanding of the dmPFC circuit’s role in social dominance. Such an expanded model could help to clarify whether the mechanisms of perseverance and aggression are distinct or overlapping in different competitive contexts.

### C. Energy cost enhances the flexibility in social competition

Neural circuits often involve tradeoffs between robustness and flexibility in various functions^27,39^. Our study has demonstrated that a circuit architecture with stronger self-excitation in PYR neurons can promote more robust effortful behaviors in social competition, endowing animals with the innate trait of perseverance, which can be advantageous for attaining dominant status. However, in such a circuit architecture, there may be a delay in initiating effortful behaviors when transitioning from the resting state of passive stillness, as depicted in Fig. 3(d). In certain contexts of social competition, animals need to react swiftly to exploit their opponents’ transient weaknesses. This raises an intriguing question: Are there any dynamic modulation mechanisms that can enhance flexibility to external signals in social competition, going beyond the circuit architecture that underlies the innate trait of perseverance?

To quantify the input-output relationship of the local dmPFC circuit, we can employ a bifurcation diagram, as illustrated in Fig. 4(a). Higher flexibility implies that even a slight change in the input can elicit a substantial shift in the system’s state, such as transitioning from the lower branch to the upper branch. Dynamically, when the system is in close proximity to the bifurcation point (indicated by dashed lines in Fig. 4(a)), an abrupt shift can occur even with a minor input alteration. However, near such a bifurcation, the system can also be driven across the boundary between the attraction basins by a small perturbation alone. In the dominance tube test, while mice may initially approach each other from opposite ends of the tube, they typically experience a brief period of stillness upon meeting before engaging in competitive behaviors^22^. In addition, the basal state of stillness is particularly evident in modified versions of the test used for specific experimental protocols, such as those involving calcium imaging^17,23^. In these cases, mice may be trained to wait at gates for short periods before the encounter begins. The robust basal state of stillness must be maintained to prevent futile assertive pushes. Therefore, even though the parameter region exhibiting bi-stability supports effortful behaviors, the dominant basal state should be passive stillness. In this regard, we argue that the initial state capable of supporting both robust basal stillness behaviors and increased flexibility to transition into effortful behaviors should be situated close to the critical location where the two alternative attractors have equal influence (indicated by the red line in Fig. 4(a)).

**FIG. 4.**
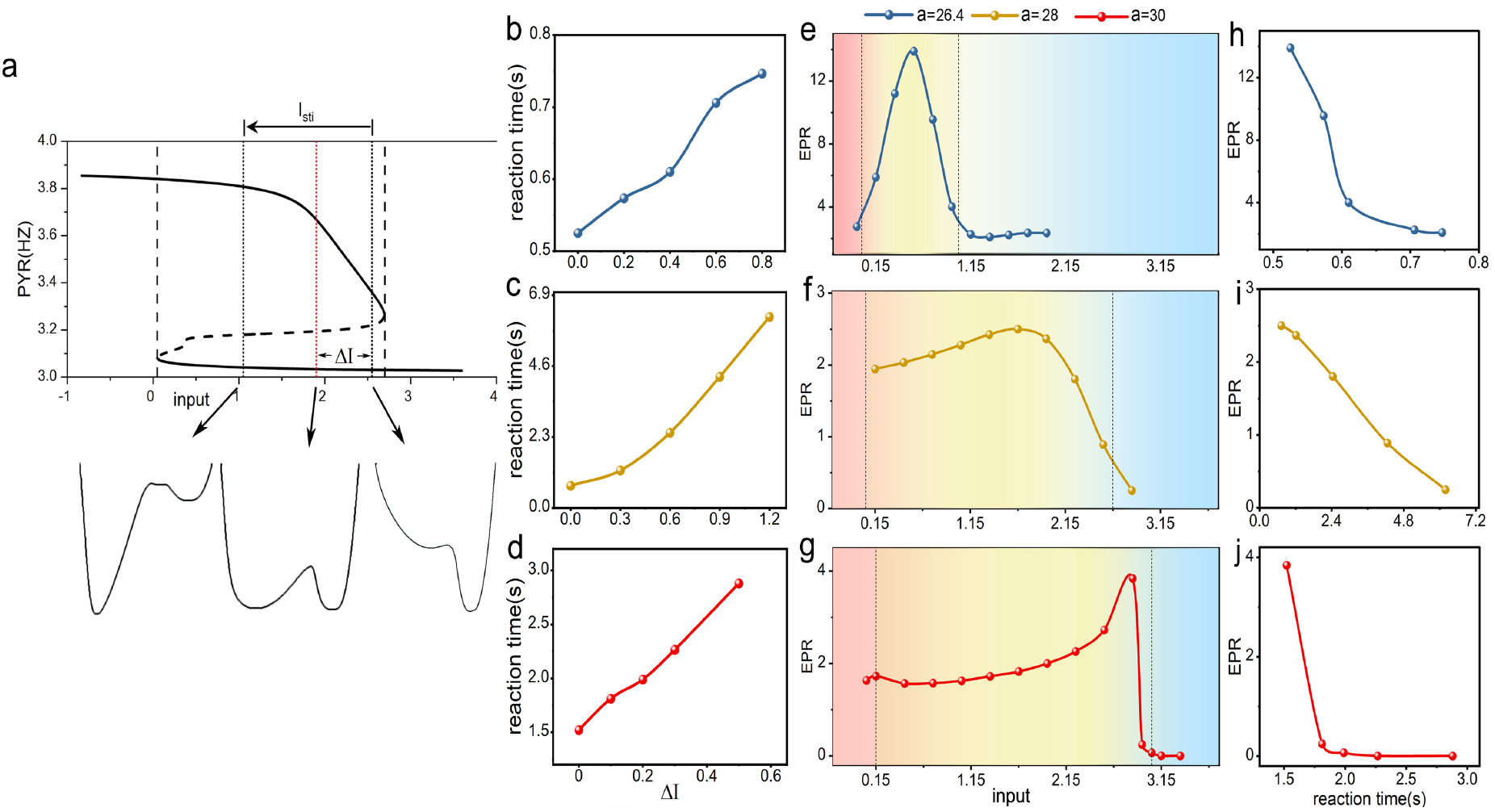
(a) The phase diagram of the model and underlying attractor landscapes for varied external inputs with the strength of self-excitation in PYR neurons *a* = 27. The total external input to PV is changed in the presence of an external stimulus input when confronting an opponent. The red line indicates the critical point where the two attractors having the same weight. (b-d) The reaction time to the external input decreases as the background input to PV neurons, which determines the basal activity of the dmPFC circuit, is close to the critical value where the two alternative behavioral states having the same weight. The variable Δ*I* measures the difference between the current value of the background input and the critical value of the background input supporting the two attractors with the same weight. The reaction time here indicates the average transition time from the passive state to the effortful state when there is a specific value in the external stimulus input to PV neurons(*I*_*sti*_ = − 0.4, − 1.5, − 2.0 for the circuit with *a* = 26.4, 28.30, respectively). (e-g) The entropy production rate(EPR), which measures the thermodynamic cost in maintaining the stable states of the non-equilibrium neural circuits, versus the varied input to PV neurons. The peak in each curve indicate the location where the two attractors having the same weight. The mono-stable regions supporting push and retreat behaviors are indicated by the red and blue colors, respectively. The bi-stable region supporting both effortful and passive behaviors is indicated by the yellow color. (h-j) The enhanced flexibility to external signals(reduced reaction time) can be achieved by larger energy cost.

The external input received by PV neurons can be divided into two components: a constant background part that determines the basal state prior to encountering an opponent, and a part associated with the external situation (e.g., the strength of the social opponent). We can refer to the latter as the external stimulus input, denoted as *I*_*sti*_. When facing an opponent, the total external input undergoes a change due to the presence of *I*_*sti*_, which is influenced by both the opponent’s state and the history of winning or losing (Fig. 4(a)). In our model, the history of winning or losing is represented by variations in the external input *I*_*sti*_ received by PV neurons when facing opponents (Fig. 4a). Animals with a history of wins would experience a larger *I*_*sti*_ compared to those with a history of losses, based on increased confidence from past successes. A larger *I*_*sti*_ then facilitates the transition from passive to effortful behaviors, recreating the “winner effect” observed experimentally^22^. As we have introduced, PV neurons receive larger external inputs when the opponent is stronger/at a higher social rank. Therefore, it is reasonable to assume that confronting weaker opponents of the same rank induces a decrease in the total input of the same magnitude (*I*_*sti*_). In addition, when facing opponents of the same rank, the magnitude of *I*_*sti*_ should be larger in circuits with stronger self-excitation in PYR neurons, corresponding to a higher social rank resulting from the trait of perseverance. This is because animals of higher rank can exhibit more confidence based on their previous winning history, leading to the delivery of a larger *I*_*sti*_ to support greater effortful behaviors such as pushes and resistance. In other words, self-confidence arising from a history of winning can facilitate the transition to effortful behaviors from the initial stillness or passive behaviors. Fig. 4(b-d) displays the average transition time from passive behaviors to effortful behaviors, also known as the reaction time, when confronted with opponents in circuits with different levels of self-excitation. It can be observed that the reaction time is shorter when the system initiates from a state closer to the location where the two attractors have equal influence (e.g., the critical point indicated by the red line in Fig. 4(a)).

An intriguing question arises: Does the enhanced flexibility come at a cost? Neural circuits constantly exchange material, information, and energy with the environment, rendering them non-equilibrium systems^36,37^. Functionally stable states in such non-equilibrium systems can only be maintained through a continuous expenditure of energy. In this study, we quantify the energy cost required to maintain stable states in the dmPFC circuits by measuring the non-equilibrium entropy production rate, as done in previous investigations^41–43^. Fig. 4(e-g) reveals a peak in the thermodynamic energy cost with varied input at the critical point. This result suggests that the reduction in reaction time and enhanced flexibility are achieved at the expense of increased energy cost (Fig. 4(h-g)). The animal, endowed with the innate trait of perseverance due to the circuit architecture with stronger self-excitation in PYR neurons, can position its basal state close to the critical point for improved flexibility to external signals, albeit at higher energy costs. This can be understood as the fact that maintaining a high level of intensity and focus to respond quickly in competitions can be exhausting for individuals.

## DISCUSSION

The tube test has long been recognized as a reliable method for assessing dominance hierarchy, and numerous studies have highlighted the role of the dorsomedial prefrontal cortex (dmPFC) in determining outcomes in social competition and dominance status^16,18–20^. The phenomenon known as the “winner effect,” a reinforcing mechanism, contributes to the establishment of dominance hierarchies, with previous winners more likely to continue winning in future contests^11–14,22^. Synaptic plasticity in the mediodorsal thalamus (MDT)-dmPFC pathway has been shown to encode this winner effect^22^. While recent research has advanced our understanding of the processing and computation of information within the dmPFC local circuit, which regulates social competition^23^, the neural mechanisms underlying the development of mental strength and complex personality traits such as courage and perseverance in social competition remain less clear.

In the context of the tube test competition, the activation or inhibition of dmPFC PYR neurons is both necessary and sufficient to induce rapid winning or losing^16^. Concurrently, PV interneurons, which project inhibitory synapses onto PYR neurons, play opposing roles in regulating competition outcomes and dominance status. Calcium imaging data from PYR and PV neurons during retreats in the tube test align with predictions based on experimental manipulations^23^. However, contrary to the expected PV neuron inhibition, an increase in PV response is observed during pushing behaviors^23^. Inspired by the well-known model of the REM sleep system, which shares similar circuit architecture with the dmPFC PV-PYR circuit^44^, we constructed a circuit model based on data obtained from previous electrophysiology recordings. By applying bifurcation analysis, we successfully reproduced the characteristic activities of different types of dmPFC neurons during various behavioral patterns observed in the tube test, including the counterintuitive excitation of PV neurons during pushes. When PV neurons receive inhibitory inputs, PYR neurons are disinhibited, and the increased activity in PYR is sustained through recurrent excitation in excitatory PYR neurons, further exciting PV neurons. The states of heightened activity in both PV and PYR correspond to the “push” behaviors exhibited in the tube test. Importantly, these distinctive patterns of dmPFC neuron activity are only observed within specific sets of control parameters that give rise to a bistable parameter region. In this bistable region, both the activated PYR and activated PV states, corresponding to effortful behaviors, and the inhibited PYR and activated PV states, corresponding to passive behaviors in the tube test, are supported. This bistability is crucial for the flexibility required in social competition, allowing for rapid transitions between effortful and passive behaviors based on the dynamics of the encounter.

To further elucidate the neural circuit mechanisms that potentially contribute to the formation of complex behavioral traits in social competitions, we quantified the attractor landscapes that govern the dynamics of the dmPFC circuit, in addition to conducting bifurcation analysis. We discovered that in a circuit architecture with stronger self-excitation in PYR neurons, states characterized by activated PYR (effortful behaviors) become significantly more robust against increased stimulus to PV neurons, which would typically induce retreats. This robustness could be interpreted as a component of what might be termed “perseverance” in a broad behavioral context. It is crucial to clarify that “perseverance” in our model refers specifically to sustained effortful behavior under external challenges, not a full evaluation of the trait as seen in psychology. Here, we focus on the neural circuit’s ability to maintain effortful behavior despite external pressures to retreat, not long-term goals, emotional regulation, or overcoming failures, which are broader aspects of human perseverance. Moreover, our findings suggest that this model-derived form of perseverance does not necessarily correlate with increased aggression, aligning with experimental observations that dominance in social hierarchies is not solely a function of aggressive behaviors^22,34,35^. This distinction is important as it underscores the non-aggressive aspects of social dominance, which can be crucial for energy conservation and strategic behavioral regulation in competitive interactions.

In our model, while the trait of perseverance (robust effortful behaviors) is enhanced by increased self-excitation in PYR neurons, this robustness comes at the cost of diminished flexibility in responding to external cues that demand immediate, forceful actions. This reduced adaptability can be detrimental in competitive situations where rapid responses are essential, such as exploiting fleeting weaknesses in opponents. Therefore, it becomes crucial to incorporate mechanisms that can dynamically adjust the balance between maintaining perseverance and enabling swift transitions to effortful actions, depending on the immediate needs of the competitive environment.

As the system approaches the bifurcation point that separates the mono-stable region, where only effortful behaviors are possible, from the bi-stable region, which supports both passive and effortful behaviors, it exhibits heightened sensitivity to external cues that could precipitate rapid transitions from passive to effortful behaviors. However, positioning the system’s resting state near this critical point can lead to instability, as minor perturbations might precipitate abrupt behavioral shifts from passivity to assertive, effortful actions. This contradicts empirical evidence suggesting that basal aggression levels do not consistently correlate with dominance status^22^. Furthermore, excessive assertive pushes can lead to an unnecessary expenditure of physical energy, which is disadvantageous in competitive scenarios. Therefore, we propose that the system’s resting state should be strategically placed within the bi-stable region surrounding the critical point, where the two stable attractors corresponding to effortful and passive behaviors have equal weighting. Here, the system can maintain stability against minor disturbances while remaining responsive to significant external stimuli. This setup allows for a balance between robustness and flexibility, essential for adaptive social behavior.

Neural networks, as typical non-equilibrium systems, are engaged in a continuous exchange of material and energy with their environment, and functional stable states in these systems are sustained through ongoing energy consumption. Our findings suggest that enhanced flexibility, which facilitates quicker reactions to external signals, incurs increased energy consumption. Minimizing energy expenditure while maximizing goal achievement is paramount for an animal’s survival. The personality trait of perseverance is instrumental in ensuring that efforts are rewarded. However, our model suggests that this trait might reduce the ability to capitalize on fleeting opportunities in competitive environments. While reduced flexibility offers the advantage of avoiding traps set by strategically superior opponents—such as baiting into a premature action—accurate assessment of situations and the selection of appropriate tactics likely involve advanced cognitive functions and cortical regions. The dynamic modulation of basal activity and resting states within our model offers a strategic choice between these alternatives. Investigating how upstream areas influence the balance between prudence and swift responsiveness represents a compelling direction for future research.

From a broader perspective, the establishment of a stable hierarchy among social animals, as facilitated by the mechanisms proposed in our model, serves to prevent unwanted conflicts among group members^19,34^, ultimately benefiting the survival of the entire population. Our findings contribute to the understanding of the neurobiological foundations for the trade-off between maintaining a stable social hierarchy and pursuing individual interests, providing insights into the complex interplay between neural circuit dynamics and social behavior.

## ACKNOWLEDGMENTS

We thank Kun Zhang and Li Xu for valuable discussions and suggestions related to this manuscript. H.Y. is grateful for support via National Natural Science Foundation of China Grants 21721003.

## DATA AVAILABILITY STATEMENT

The data that support the findings of this study are available from the corresponding author upon reasonable request.

## Appendix A: Circuit model in controlling social competition

Applying the dominance tube test, researchers have identified the dorsomedial prefrontal cortex (dmPFC) neural population playing a crucial role in controlling social competition. In the tube test competition, the activation of the dmPFC PYR neurons is both necessary and sufficient to quickly induce winning. Activation/inhibition of PV interneurons in dmPFC, which inhibit PYR neurons, induces losing/wining.

The working model of the local PYR-PV circuit in dmPFC was constructed based on experimental recordings from several key studies in the field. We primarily drew upon the work of Zhang et al.^23^, which provided valuable insights into the circuit architecture involving reciprocal connections between PYR and PV neurons and their corresponding dynamics during different behavioral epochs in the dominance tube test. Specifically, Zhang et al. observed that PYR neurons are activated (increased calcium signals) during push epochs and inhibited (decreased calcium signals) during retreat epochs, while PV neurons show increased activity during both push and retreat epochs. To further refine our model, we incorporated mean firing rate data from Zhou et al.^22^. We used approximate mean firing rates of PYR and PV neurons during different behavioral states (e.g., push, retreat, resistance) to calibrate our model. For example, PYR neurons exhibited a baseline mean firing rate of approximately 3 −3.5 Hz, increasing to about 4 Hz during push behaviors, while PV neurons showed baseline firing rates of approximately 3.5 Hz, increasing to about 7 Hz during retreat behaviors. These experimental observations guided our parameter tuning process, allowing us to adjust our model to reproduce key features of PYR and PV neuron dynamics in the dmPFC during social competition behaviors in a physiologically plausible manner.

Based on these experimental recordings, a working model of the local PYR-PV circuit in dmPFC can be constructed. A shown in Fig.1, PYR neurons excite themselves and also PV neurons. PV neurons, conversely, inhibit the activity of PYR neurons. Inspired by the famous model of the REM sleep system which shares the similar circuit architecture with the dmPFC PV-PYR circuit, we introduced nonlinear terms describing the inner and inter-actions of PYR and PV neurons. The dynamics of two populations can be described by the following equations:

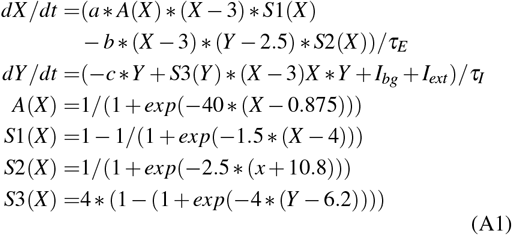

*X* and *Y* here are the average activities of excitatory pyramidal (PYR) neurons and parvalbumin(PV) interneurons. The term representing the excitatory feedback in PYR neurons consists of two parts: a constant part indicated by the control parameter *a* and a normalized nonlinear function *A*(*X*) that depends on the mean activity of PYR neural population. *S*1(*X*) and *S*2(*Y*) are saturation functions constraining maximal firing rates in PYR(X) and PV(Y) populations. The term *S*2(*X*) limits the inhibition to PYR from PV for low activity in PYR. The standard set of parameters for this model is as follows: *a* = 27.2, *b* = 7.2, *c* = 1.

While direct experimental quantification of the specific circuit parameters is still lacking, the values chosen aim to reproduce key features observed experimentally in a physiologically plausible manner^23^. In future work, incorporating stochasticity in a more biophysically realistic way and tuning the model to experimentally measured variability could allow indirect empirical constraints on the parameters. Ultimately, targeted electrophysiology is needed to directly measure parameters like connection strengths. The decay time constants for the excitatory/inhibitory neurons whose synaptic transmissions mediated by NMDA/GABA receptors are set as: *τ*_*E*_ = 100*ms, τ*_*I*_ = 2*ms*.

The dynamics of this circuit model that controls behavioral outputs in social competition are determined by both the circuit achitecture(measured by the connection strengths *a*, etc) and the inputs to PV neurons including a constant part from the background *I*_*bg*_, which determines the resting state before confronting an opponent and a part associated with the external situation *I*_*sti*_ (e.g., is the social opponent strong or weak).

In the model, we introduced a self-excitation mechanism in the PYR population, which serves as an abstraction for the recurrent connectivity and potential neuromodulatory feedback that PYR neurons can receive in vivo. This self-excitation is depicted as a function of the mean activity level of PYR neurons, reflecting the capacity of these neurons to sustain or amplify their activity through intrinsic properties or through network interactions with other neurons within the dmPFC or from afferent terminals.

To bridge the gap between our computational model and biological reality, it is important to consider that self-excitation in live animals is likely to be mediated by a combination of factors. These may include direct recurrent connections between PYR neurons, which have been observed in the prefrontal cortex, as well as indirect pathways involving other interneurons or long-range inputs that could facilitate a feedback loop leading to sustained PYR neuron activity. More- over, neuromodulatory systems, such as dopaminergic or cholinergic inputs, might also contribute to the self-excitation by modulating the excitability of PYR neurons. Our model simplifies these complex interactions into a single parameter for computational tractability, but we recognize that the underlying biological mechanisms are multifaceted and extend beyond the scope of this model.

## Appendix B: Non-equilibrium landscape and flux framework for general neural circuits

The concept of an attractor landscape is utilized to illustrate the correspondence between neural activity patterns and various cognitive states, with each state depicted as a basin within this landscape. This analogy is prevalent in discussions on cognitive functions such as associative memory retrieval and classification^45,46^. However, the mathematical representation and computation of this concept, particularly in terms of stability and transition rates, are confined to models with specific symmetry assumptions, such as those seen in the symmetrical neural circuit of the original Hopfield model^45,46^. To extend the understanding of global stability of states associated with cognitive functions, we have developed a non-equilibrium potential landscape and flux theory that applies to a broader spectrum of neural network models, including those without specific symmetries^36^.

For complex systems composed of numerous interacting components, tracking the microscopic trajectories of each component through solving corresponding dynamical equations is challenging. Yet, the macroscopic emergent behaviors of the system can be approximated by the dynamical evolution of macroscopic observables, influenced by deterministic and stochastic forces. An exemplary model of such dynamics is the stochastic Langevin particle dynamics. We consider a system governed by Langevin equations: 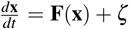, where the state variables are denoted by the vector **x**, the driving force by the vector **F**(**x**), and Gaussian white noise by *ζ*, with an ensemble average of zero. This indicates that the average noise over multiple realizations is zero. The autocorrelations of the noise are described by *< ζ* (**x**, *t*)*ζ* (**x**, *t*^′^) *>*= 2*D***D**(**x**)*δ* (*t* −*t*^′^), where *D* is a scale factor and **D**(**x**) is the diffusion tensor. These variables evolve according to nonlinear dynamical laws due to complex system interactions, leading to unpredictable behaviors when coupled with stochastic influences.

The stochastic differential equations govern the temporal trajectories of neural firing, with their probabilistic evolution characterized by the Fokker-Planck equation for the probability density, as referenced in several studies^36,47,48^:

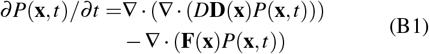

The corresponding Fokker-Planck equation can be expressed as follows:

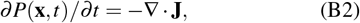

where the probability flux is defined by:

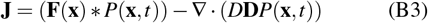

The alteration in local probability *P*(**x**, *t*) over time is governed by the net probability flux **J**, which ensures local conservation of probability. For *t* → ∞, the divergence of the flux becomes zero, indicating a steady state:

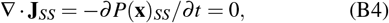

Here, **J**_*SS*_ denotes the steady-state flux. However, this condition of zero divergence does not necessitate the flux itself being zero. A non-zero flux indicates a directional flow of probabilities, disrupting the detailed balance and signifying operation out of thermodynamic equilibrium. Moreover, a divergence-free flux implies the absence of sources or sinks for the probability to go to or come from, and hence, locally, the flux must be rotational.

Upon reaching a steady state, the steady-state probability distributions can be solved through the steady-state equation:

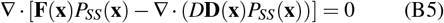

Additionally, the non-equilibrium potential function *U* = −ln *P*_*SS*_(**x**) is defined, analogous to the Boltzmann law in equilibrium statistical mechanics. The driving forces are decomposed as follows^36,37,47–49^:

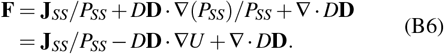

Contrary to equilibrium systems, where dynamics are solely influenced by the gradient of the underlying energy function resulting in zero net flux, non-equilibrium systems are influenced by both the potential gradient and the curl flux (**J**_*SS*_). The non-equilibrium potential *U* is instrumental in characterizing the global behavior of such systems.

In a steady state, non-equilibrium thermodynamic systems are characterized by continuous entropy production^36,37,49^. Although the entropy of a non-equilibrium system in a steady state remains constant over time, there is an equivalent flow of entropy to the surroundings that matches the entropy generated internally. To compute the entropy production, it is essential to focus on the entropy associated with the time-dependent probability distribution. Utilizing the Fokker-Planck equation and the definition of probability flux **J**(**x**), the time derivative of the system entropy *dS/dt* can be articulated as a sum of two terms:

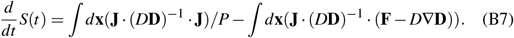

The first term on the right-hand side of the equation 8 is always non-negative due to the positive definite nature of the diffusion matrix **D**, representing the total entropy production rate, which is the global thermodynamic dissipation or cost directly linked to the non-equilibrium flux^37^:

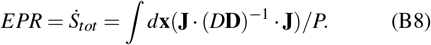

where EPR stands for entropy production rate. The second term on the right-hand side of equation 8 can then be regarded as the entropy flux from the system to the environment:

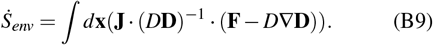

The total entropy change is equal to the sum of the entropy change of the system and that of the environment, leading to the emergence of the generalized non-equilibrium first law of thermodynamics:

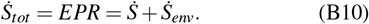

The entropy change of the non-equilibrium system 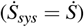 can either increase or decrease due to the entropy flow to the environments, while the total entropy change of the system and the environments, represented by the entropy production rate, is always non-negative. This leads to the generalized non-equilibrium thermodynamic second law:

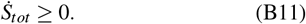

## Appendix C: Calculation of transition time and barrier height

To further analyze the dynamics of our model, we quantify two key measures: transition time and barrier height. These metrics provide insights into the stability of attractor states and the difficulty of transitions between them, which are crucial for understanding the robustness of effortful and passive behaviors in our model of social competition.

Transition time: In our model, the transition time refers to the average time it takes for the system to switch from one attractor state to another in the bistable region. We calculate this using the following method: The dynamical trajectories of the system are computed using the Runge-Kutta method. Random fluctuations are introduced through a noise term, implemented as uncorrelated standard Gaussian noise with zero mean and a variance of 0.02. For each data point, we conduct over 10,000 trials to obtain a statistically robust average transition time.

Barrier height: The barrier height represents the potential difference between an attractor state and the saddle point separating it from another attractor state. We calculate this using the potential landscape derived from the steady-state probability distribution: We define the potential landscape as *U* = −ln *P*_*SS*_(**x**), where *P*_*SS*_(**x**) is the steady-state probability distribution. The barrier height is then computed as: Barrier Height = *U*_*saddle*_ −*U*_*min*_, where *U*_*min*_ is the potential minimum of one local attractor, and *U*_*saddle*_ is the potential at the saddle point between two attractors.

These calculations allow us to quantify the stability of the attractor states and the difficulty of transitions between them, providing insights into the robustness of effortful and passive behaviors in our model of social competition.

## Appendix D: Quantification of neural circuit energy expenditure

Biological systems, including neural circuits, expend energy to facilitate various essential functions^49,50^. Although increased neuronal firing rates typically correlate with heightened energy requirements^51^, using firing rate as the sole metric to quantify energy expenditure in neural circuits does not encompass significant aspects of the brain’s total energy use. As elucidated by Marcus E. Raichle through the “dark energy” concept, a substantial portion of energy is allocated to baseline cellular functions that occur independently of external stimuli, with task-related energy use potentially representing only 0.5 −1.0% of the total consumption. Moreover, approximately one-third of energy supports “housekeeping” activities such as protein and lipid synthesis, proton leakage across the mitochondrial membrane, and cytoskeletal modifications. These ongoing, non-signaling processes account for 25 −50% of the brain’s energy expenditure^52^, indicating that firing rates alone are insufficient for measuring the full spectrum of energy costs associated with neural circuits.

In our research, we utilize the entropy production rate (EPR) to quantitatively assess the energy expenditure of biological systems, specifically neural circuits, by examining their continuous energy dissipation. Our approach is based on a solid theoretical foundation, bolstered by a series of both theoretical and empirical studies^37,41–43,53,54^, which establish EPR as a crucial thermodynamic quantity for quantifying the ongoing energy dissipation inherent in biological systems.

To implement this concept, we view the biological system as an open system maintained far from equilibrium through the continuous input and dissipation of energy, primarily via the hydrolysis of ATP and GTP molecules. The overall thermodynamic cost includes both the free energy input, or known as the house-keeping heat *Q*_*HK*_^55^, required to sustain the non-equilibrium steady state and the generalized free energy relaxation 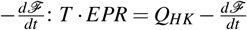. The total thermodynamic dissipation or cost is quantified as the product of EPR and temperature, covering both steady-state maintenance and free energy transformations^37,41,53^.

The hydrolysis of ATP, a primary source of entropy production, supplies the free energy necessary for driving endergonic biochemical reactions that maintain non-equilibrium conditions in living cells. The ATP/ADP ratio, which reflects these conditions, results in a chemical potential difference Δ*G* ∼ ln(*ATP/ADP*) analogous to the voltage in electric circuits, acting as the system’s energy pump^56^. As demonstrated, the EPR is directly related to the non-equilibrium flux, linking it to the chemical reaction flux^56^. The rate of ATP/GTP hydrolysis and the consumption of other metabolic fuels are thus directly associated with the EPR, quantifying the minimal rate of heat dissipation needed to counteract fluctuation-driven disorder. A higher EPR indicates increased expenditure and metabolic demand required to sustain the dynamics of a non-equilibrium living system.

The significance of EPR as a measure of energy cost is empirically validated by studies on the Escherichia coli chemosensory system^43^. In this critical research, the entropy production rate was used to quantify energy consumption, revealing an intrinsic energy-speed-accuracy relationship. The study demonstrated that the non-equilibrium state of the system’s negative feedback mechanism necessitates energy dissipation for precise sensory adaptation. The experimental validation of theoretical predictions in E. coli chemotaxis confirms the utility of EPR in quantifying energy costs in biological adaptation mechanisms.

Our application of EPR to neural circuits is informed by these insights, providing a comprehensive framework for evaluating continuous and non-equilibrium energy demands. This methodological advancement enables us to measure energy dissipation beyond transient neuronal firing, offering a more inclusive perspective on the brain’s energy utilization.

## References

1 E. O. Wilson, Sociobiology: The new synthesis (Harvard University Press, 2000).

2 B. J. Cole, “Dominance hierarchies in leptothorax ants,” Science 212, 83– 84 (1981).

3 R. Dunbar and E. Dunbar, “Dominance and reproductive success among female gelada baboons,” Nature 266, 351–352 (1977).

4 L. Grosenick, T. S. Clement, and R. D. Fernald, “Fish can infer social rank by observation alone,” Nature 445, 429–432 (2007).

5 F. B. Bercovitch and A. S. Clarke, “Dominance rank, cortisol concentrations, and reproductive maturation in male rhesus macaques,” Physiology & behavior 58, 215–221 (1995).

6 R. M. Sapolsky, “The influence of social hierarchy on primate health,” science 308, 648–652 (2005).

7 N. So, B. Franks, S. Lim, and J. P. Curley, “A social network approach reveals associations between mouse social dominance and brain gene expression,” PloS one 10, e0134509 (2015).

8 T. Zhou, C. Sandi, and H. Hu, “Advances in understanding neural mechanisms of social dominance,” Current Opinion in Neurobiology 49, 99–107 (2018).

9 S. J. Mooney, D. E. Peragine, G. A. Hathaway, and M. M. Holmes, “A game of thrones: Neural plasticity in mammalian social hierarchies,” Social neuroscience 9, 108–117 (2014).

10 R. D. Fernald, “Cognitive skills needed for social hierarchies,” in Cold Spring Harbor symposia on quantitative biology, Vol. 79 (Cold Spring Harbor Laboratory Press, 2014) pp. 229–236.

11 E. Bonabeau, G. Theraulaz, and J.-L. Deneubourg, “Dominance orders in animal societies: the self-organization hypothesis revisited,” Bulletin of mathematical biology 61, 727–757 (1999).

12 K. Kura, M. Broom, and A. Kandler, “A game-theoretical winner and loser model of dominance hierarchy formation,” Bulletin of mathematical biology 78, 1259–1290 (2016).

13 M.-Y. Chou, R. Amo, M. Kinoshita, B.-W. Cherng, H. Shimazaki, M. Agetsuma, T. Shiraki, T. Aoki, M. Takahoko, and M. Yamazaki, “Social conflict resolution regulated by two dorsal habenular subregions in zebrafish,” Science 352, 87–90 (2016).

14 L. A. Dugatkin and M. Druen, “The social implications of winner and loser effects,” Proceedings of the Royal Society of London. Series B: Biological Sciences 271, S488–S489 (2004).

15 G. Lindzey, H. Winston, and M. Manosevitz, “Social dominance in inbred mouse strains,” Nature 191, 474–476 (1961).

16 F. Wang, J. Zhu, H. Zhu, Q. Zhang, Z. Lin, and H. Hu, “Bidirectional control of social hierarchy by synaptic efficacy in medial prefrontal cortex,” Science 334, 693–697 (2011).

17 Z. Fan, H. Zhu, T. Zhou, S. Wang, Y. Wu, and H. Hu, “Using the tube test to measure social hierarchy in mice,” Nature Protocols 14, 819–831 (2019).

18 M. F. Rushworth, R. B. Mars, and J. Sallet, “Are there specialized circuits for social cognition and are they unique to humans?” Current opinion in neurobiology 23, 436–442 (2013).

19 F. Wang, H. W. Kessels, and H. Hu, “The mouse that roared: neural mechanisms of social hierarchy,” Trends in neurosciences 37, 674–682 (2014).

20 H. Zhu and H. Hu, “Brain¡^-^s neural switch for social dominance in animals,” Sci China Life Sci 61, 113–114 (2018).

21 T. B. Franklin, B. A. Silva, Z. Perova, L. Marrone, M. E. Masferrer, Y. Zhan, A. Kaplan, L. Greetham, V. Verrechia, and A. Halman, “Prefrontal cortical control of a brainstem social behavior circuit,” Nature neuroscience 20, 260–270 (2017).

22 T. Zhou, H. Zhu, Z. Fan, F. Wang, Y. Chen, H. Liang, Z. Yang, L. Zhang, L. Lin, and Y. Zhan, “History of winning remodels thalamo-pfc circuit to reinforce social dominance,” Science 357, 162–168 (2017).

23 C. Zhang, H. Zhu, Z. Ni, Q. Xin, T. Zhou, R. Wu, G. Gao, Z. Gao, H. Ma, and H. Li, “Dynamics of a disinhibitory prefrontal microcircuit in controlling social competition,” Neuron 110, 516–531. e6 (2022).

24 B. S. Porter, K. L. Hillman, and D. K. Bilkey, “Anterior cingulate cortex encoding of effortful behavior,” Journal of neurophysiology 121, 701–714 (2019).

25 M. E. Walton, D. M. Bannerman, and M. F. Rushworth, “The role of rat medial frontal cortex in effort-based decision making,” Journal of Neuroscience 22, 10996–11003 (2002).

26 K. L. Hillman, “Cost-benefit analysis: the first real rule of fight club?” Frontiers in neuroscience 7, 248 (2013).

27 M. S. Goldman, J. Golowasch, E. Marder, and L. Abbott, “Global structure, robustness, and modulation of neuronal models,” Journal of Neuroscience 21, 5229–5238 (2001).

28 W. R. Softky and C. Koch, “The highly irregular firing of cortical cells is inconsistent with temporal integration of random epsps,” Journal of Neuroscience 13, 334–350 (1993).

29 M. N. Shadlen and W. T. Newsome, “The variable discharge of cortical neurons: implications for connectivity, computation, and information coding,” Journal of neuroscience 18, 3870–3896 (1998).

30 C. Hanus and E. M. Schuman, “Proteostasis in complex dendrites,” Nature Reviews Neuroscience 14, 638 (2013).

31 D. Kvitsiani, S. Ranade, B. Hangya, H. Taniguchi, J. Huang, and A. Kepecs, “Distinct behavioural and network correlates of two interneuron types in prefrontal cortex,” Nature 498, 363–366 (2013).

32 H.-J. Pi, B. Hangya, D. Kvitsiani, J. I. Sanders, Z. J. Huang, and A. Kepecs, “Cortical interneurons that specialize in disinhibitory control,” Nature 503, 521–524 (2013).

33 A. Kepecs and G. Fishell, “Interneuron cell types are fit to function,” Nature 505, 318–326 (2014).

34 R. C. Francis, “The effects of bidirectional selection for social dominance on agonistic behavior and sex ratios in the paradise fish (macropodus opercularis),” Behaviour 90, 25–44 (1984).

35 D. Benton, J. C. Dalrymple-Alford, and P. F. Brain, “Comparisons of measures of dominance in the laboratory mouse,” Animal Behaviour 28, 1274– 1279 (1980).

36 H. Yan, L. Zhao, L. Hu, X. D. Wang, E. K. Wang, and J. Wang, “Nonequilibrium landscape theory of neural networks,” Proceedings of the National Academy of Sciences of the United States of America 110, E4185–E4194 (2013).

37 J. Wang, “Landscape and flux theory of non-equilibrium dynamical systems with application to biology,” Advances in Physics 64, 1–137 (2015).

38 H. Yan, B. Li, and J. Wang, “Non-equilibrium landscape and flux reveal how the central amygdala circuit gates passive and active defensive responses,” Journal of the Royal Society Interface 16 (2019), Artn 2018075610. 1098/Rsif.2018.0756.

39 H. Yan and J. Wang, “Non-equilibrium landscape and flux reveal the stability-flexibility-energy tradeoff in working memory,” PLoS Computational Biology 16, e1008209 (2020).

40 C. M. Williamson, I. S. Klein, W. Lee, and J. P. Curley, “Immediate early gene activation throughout the brain is associated with dynamic changes in social context,” Social Neuroscience 14, 253–265 (2019).

41 H. Ge and H. Qian, “Dissipation, generalized free energy, and a self-consistent nonequilibrium thermodynamics of chemically driven open subsystems,” Physical Review E 87, 062125 (2013).

42 Y. Cao, H. Wang, Q. Ouyang, and Y. Tu, “The free-energy cost of accurate biochemical oscillations,” Nature physics 11, 772 (2015).

43 G. Lan, P. Sartori, S. Neumann, V. Sourjik, and Y. Tu, “The energy^..^cspeed^..^caccuracy trade-off in sensory adaptation,” Nature physics 8, 422 (2012).

44 R. McCarley and S. Massaquoi, “Neurobiological structure of the revised limit cycle reciprocal interaction model of rem cycle control,” Journal of sleep research 1, 132–137 (1992).

45 J. J. Hopfield, “Neural networks and physical systems with emergent collective computational abilities,” Proceedings of the National Academy of Sciences of the United States of America-Biological Sciences 79, 2554– 2558 (1982).

46 J. J. Hopfield and D. W. Tank, “Computing with neural circuits - a model,” Science 233, 625–633 (1986).

47 J. Wang, L. Xu, and E. K. Wang, “Potential landscape and flux framework of nonequilibrium networks: Robustness, dissipation, and coherence of biochemical oscillations,” Proceedings of the National Academy of Sciences of the United States of America 105, 12271–12276 (2008).

48 J. Wang, K. Zhang, L. Xu, and E. Wang, “Quantifying the waddington landscape and biological paths for development and differentiation,” Proceedings of the National Academy of Sciences of the United States of America 108, 8257–8262 (2011).

49 H. Yan, K. Zhang, and J. Wang, “Physical mechanism of mind changes and tradeoffs among speed, accuracy, and energy cost in brain decision making: Landscape, flux, and path perspectives,” Chinese Physics B 25 (2016).

50 E. Eisenberg and T. L. Hill, “Muscle contraction and free energy transduction in biological systems,” Science 227, 999–1006 (1985).

51 D.-P. Yang, H.-J. Zhou, and C. Zhou, “Co-emergence of multi-scale cortical activities of irregular firing, oscillations and avalanches achieves cost-efficient information capacity,” PLOS Computational Biology 13, e1005384 (2017).

52 E. Engl and D. Attwell, “Non-signalling energy use in the brain,” The Journal of Physiology 593, 3417–3429 (2015).

53 H. Ge and H. Qian, “Physical origins of entropy production, free energy dissipation, and their mathematical representations,” Physical Review E 81, 051133 (2010).

54 X. Yang, M. Heinemann, J. Howard, G. Huber, S. Iyer-Biswas, G. Le Treut, M. Lynch, K. L. Montooth, D. J. Needleman, S. Pigolotti, J. Rodenfels, P. Ronceray, S. Shankar, I. Tavassoly, S. Thutupalli, D. V. Titov, J. Wang, and P. J. Foster, “Physical bioenergetics: Energy fluxes, budgets, and constraints in cells,” Proceedings of the National Academy of Sciences 118, e2026786118 (2021).

55 T. Hatano and S.-i. Sasa, “Steady-state thermodynamics of langevin systems,” Physical Review Letters 86, 3463–3466 (2001).

56 L. Xu, H. Shi, H. Feng, and J. Wang, “The energy pump and the origin of the non-equilibrium flux of the dynamical systems and the networks,” The Journal of chemical physics 136, 04B621 (2012).

